# Warm temperature suppresses plant systemic acquired resistance by intercepting *N*-hydroxypipecolic acid biosynthesis

**DOI:** 10.1101/2023.10.31.564368

**Authors:** Alyssa Shields, Lingya Yao, Christina A.M. Rossi, Paula Collado Cordon, Jong Hum Kim, Wasan Mudher Abu AlTemen, Sha Li, Eric J. R. Marchetta, Vanessa Shivnauth, Tao Chen, Sheng Yang He, Xiu-Fang Xin, Christian Danve M. Castroverde

## Abstract

Climate warming influences disease development by targeting critical components of the plant immune system, including pattern-triggered immunity (PTI), effector-triggered immunity (ETI) and production of the central defence hormone salicylic acid (SA) at the primary pathogen infection site. However, it is not clear if and/or how temperature impacts systemic immunity. Here we show that pathogen-triggered systemic acquired resistance (SAR) in *Arabidopsis thaliana* is suppressed at elevated temperature. This was accompanied by global downregulation of SAR-induced genes at elevated temperature. Abolished SAR under warmer conditions was associated with reduced biosynthesis of the SAR metabolite *N*-hydroxypipecolic acid (NHP) in *Arabidopsis* and other plant species (such as tomato and rapeseed), as demonstrated by downregulation of NHP biosynthetic genes (*ALD1* and *FMO1*) and reduced NHP and pipecolic acid (Pip) levels. Although multiple SAR signals have been shown previously, exogenous NHP or Pip was sufficient to restore disease protection at elevated temperature, indicating that heat-mediated SAR suppression is due to downregulation of the NHP pathway. Along with *ALD1* and *FMO1*, local and systemic expression of the SA biosynthetic gene *ICS1* were also suppressed at warm temperature. Finally, we define a transcriptional network controlling thermosensitive NHP pathway via the master transcription factors CBP60g and SARD1. Our findings demonstrate that warm temperatures impact not only local but also systemic immunity by impinging on NHP biosynthesis, providing a roadmap towards engineering climate-resilient plant immune systems.

## Introduction

Warming global temperatures due to climate change pose serious threats to the natural environment and human civilization (Altizer et al., 2013; Deutsch et al., 2018; Cavicchioli et al., 2019; Delgado-Baquerizo et al., 2020). In agricultural systems, increasing average temperatures are predicted to significantly reduce yields of the world’s major crops (Zhao et al., 2017; Chaloner et al., 2021; Singh et al., 2023). As many crop varieties are bred to be grown outside the native growth ranges of their wild relatives, it is important to understand the impact of elevated temperatures on plant growth, development, immunity and phenology. Together with other environmental factors, temperature profoundly affects various aspects of plant physiology, ranging from growth and development (Quint et al., 2016; Lippman et al., 2019; Hayes et al., 2020; Vu et al., 2021; Castroverde and Dina, 2021) to stress responses and immunity (Velasquez et al., 2018; Kim et al., 2021; Castroverde and Dina, 2021; Singh et al., 2023). Environmentally compromised immune signaling is associated with increased plant disease development, as postulated as the plant disease triangle (Stevens, 1960; Colhoun, 1973; Roussin-Léveillée et al., 2024).

Elevated temperatures target various components of the plant immune system (Velasquez et al., 2018; Cheng et al., 2019; Cohen and Leach, 2020; Kim et al., 2021; Castroverde et al., 2024), which encompass both cell surface (transmembrane) and intracellular immune receptors (Jones and Dangl, 2006; Zhou and Zhang, 2020; Kim and Castroverde, 2020; Ngou et al., 2022). Cell surface immune receptors recognize apoplastic immunogenic molecules, typically conserved pathogen-associated molecular patterns (PAMPs; Bigeard et al., 2015; Gust et al., 2017; DeFalco and Zipfel, 2021; Shu et al., 2023), while intracellular nucleotide-binding leucine-rich receptors (NLRs) perceive pathogen effectors and/or effector-induced host modifications (Cui et al., 2015; Jones et al., 2016; Saur et al., 2021; Contreras et al., 2023). There is evidence that warm temperatures lead to reduced abundance, membrane localization or signaling of cell surface receptors (De Jong et al., 2002; Janda et al., 2019; Hilleary et al., 2020) and perturbed nuclear localization or activation of certain NLRs (Zhu et al., 2010; Mang et al., 2012; Cheng et al., 2013; Venkatesh and Kang, 2019).

Recent studies show that both PRR signaling and NLR signaling activate convergent downstream signaling, including increased cytoplasmic calcium influx, oxidative burst, phosphorylation cascades and production of defence hormone salicylic acid (SA), leading to overlapping defence gene expression and metabolism at the primary infection site (Pieterse et al., 2009; Ngou et al., 2021; Yuan et al., 2021). Previous research has shown that elevated temperature suppresses pathogen-induced biosynthesis of the central defence hormone salicylic acid (SA) (Malamy et al., 1992; Huot et al., 2017; Li et al., 2020; Kim et al., 2022; Rossi et al., 2023), which aligns with temperature modulation of other plant hormones (Franklin et al., 2011; Sun et al., 2012; Ibañez et al., 2018; Martínez et al., 2018; Ferrero et al., 2019; Bruessow et al., 2021; Castroverde and Dina, 2021). Apart from resistance at the local infection site, SA is also involved in systemic immunity known as systemic acquired resistance (SAR) (Fu and Dong, 2013; Zhang and Li, 2019; Kachroo and Kachroo, 2020; Lim et al., 2020; Peng et al., 2021). Primary infection and immune activation lead to SAR, which primes non-infected systemic tissues against secondary infections (Cameron et al., 1994; Maldonado et al., 2002; Fu and Dong, 2013; Yu et al., 2013; Vlot et al., 2021; Zeier, 2021). SAR is widely conserved across the plant kingdom, leading to broad-spectrum host defences by rapidly generating mobile signals at the infection site and transporting them throughout the plant (Carella et al., 2016; Chen et al., 2018; Hartmann et al., 2018; Kachroo and Kachroo, 2020; Shine et al., 2022; Cao et al., 2024).

A major SAR signal is the immune-activating metabolite *N*-hydroxypipecolic acid (NHP) (Chen et al., 2018; Hartmann and Zeier, 2018; Hartmann et al., 2018; Yildiz et al., 2021; Löwe et al., 2023). NHP has been shown to induce SAR, and blocking its biosynthesis results in SAR abolishment (Song et al., 2004; Mishina and Zeier, 2006; Jing et al., 2011; Návarová et al., 2012; Huang et al., 2020; Brambilla et al., 2023). NHP biosynthesis follows three consecutive enzymatic reactions (Hartmann and Zeier, 2018), involving the aminotransferase AGD2-LIKE DEFENSE RESPONSE PROTEIN 1 (ALD1; Zeier, 2013; Bernsdorff et al., 2016), the reductase SAR-DEFICIENT 4 (SARD4; Ding et al., 2016; Hartmann et al., 2017) and *N*-hydroxylase FLAVIN-DEPENDENT MONOOXYGENASE 1 (FMO1), which converts pipecolic acid (Pip) to *N*-hydroxypipecolic acid (NHP; Chen et al., 2018; Hartmann et al., 2018). *ALD1*, *SARD4* and *FMO1* gene expression and NHP levels are induced systemically following pathogenic attack at a primary site (Song et al., 2004; Mishina and Zeier, 2006; Ding et al., 2016; Bernsdorff et al., 2016; Hartmann et al., 2017; Yildiz et al., 2021).

Currently, the impact of elevated temperature on plant systemic immunity is underexplored. Although previous studies have shown that increased temperatures have a negative effect on several aspects of the plant immune system that occur in the local/primary sites of infection (like PTI, ETI and SA production), it is unknown if and how elevated temperature affect NHP-mediated plant immunity and the SAR response. In our study, we show that *e*levated temperature suppresses SAR-associated NHP biosynthetic gene expression and production of SAR signals Pip and NHP in *Arabidopsis* and other plant species. Because it has been shown that the NHP immune pathway is redundantly controlled by the master transcription factors CALMODULIN-BINDING PROTEIN 60-LIKE G (CBP60g) and SAR-DEFICIENT 1 (SARD1) (Sun et al., 2015; Sun et al., 2018; Huang et al., 2020), we further found that warm temperature-suppressed plant systemic immunity is restored by constitutive *CBP60g/SARD1* gene expression or exogenous Pip or NHP application. Collectively, our study indicates that CBP60g and SARD1 control the temperature-vulnerability of *Arabidopsis* systemic immunity by regulating Pip/NHP biosynthesis.

## Results

### Warm temperature affects systemic acquired resistance in *Arabidopsis* plants

To determine if SAR is affected by temperature, lower leaves of *Arabidopsis* Col-0 plants were inoculated with the virulent bacterial pathogen *Pst* DC3000 as the primary challenge. Two days after primary infection, upper systemic leaves were inoculated with the pathogen *Pst* DC3000 as the secondary challenge. As shown in Figure 1A, primary pathogen infection expectedly lowered the systemic bacterial counts after secondary pathogen challenge at 23°C but not at 28°C in Col-0 plants. This indicates that *Arabidopsis* SAR due to local *Pst* DC3000 infection is effective at normal but not at elevated temperature, suggesting that temperature regulates plant systemic immunity.

**Figure 1.**
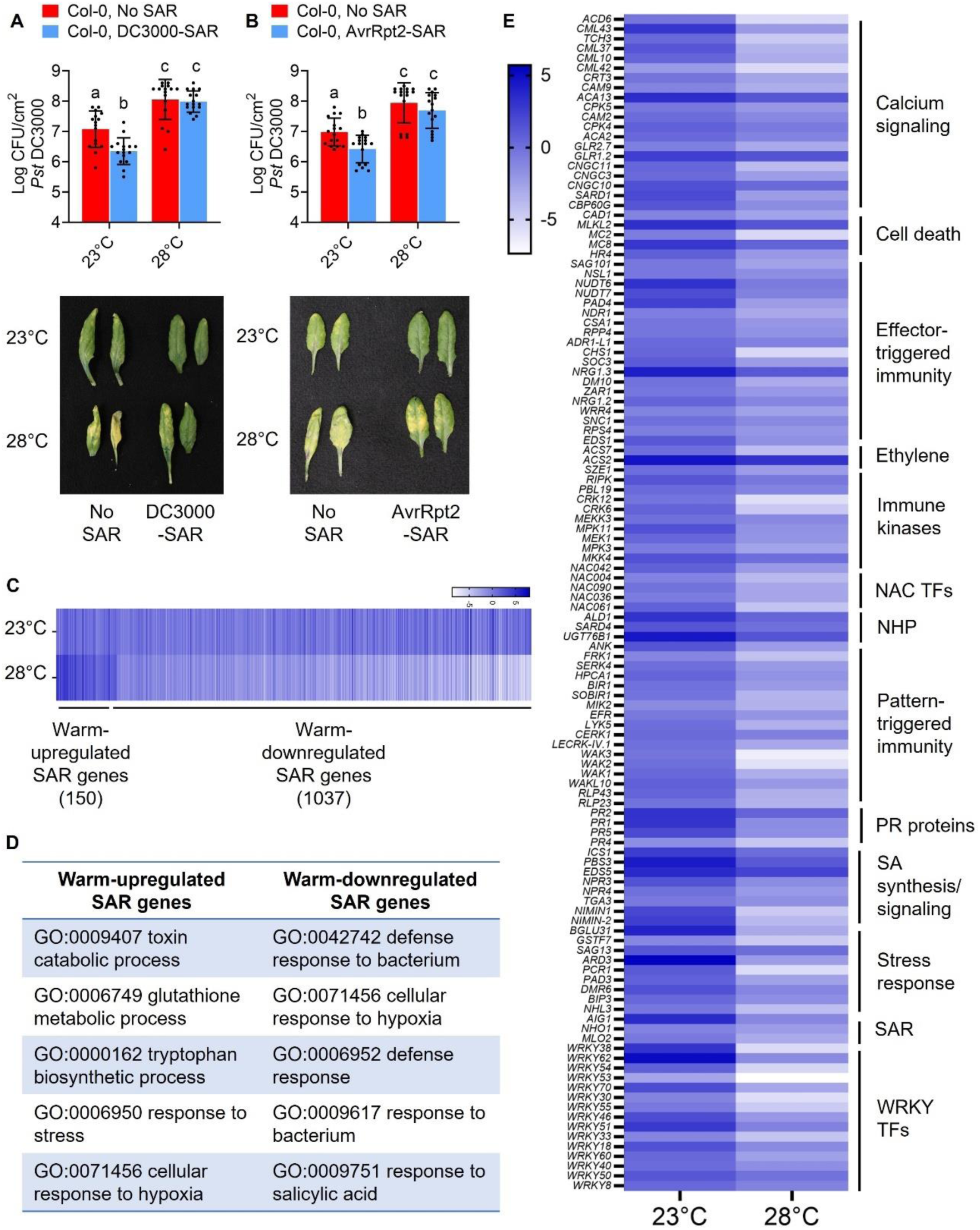
*Arabidopsis* SAR is suppressed at elevated temperature. (A, B) Lower leaves of four-week-old *Arabidopsis* Col-0 plants were infiltrated with 0.25 mM MgCl_2_ (mock) and *Pst* DC3000 (OD600 = 0.02) in (A) or *Pst* DC3000/AvrRpt2 (OD600 = 0.02) in (B). Plants were then incubated at either 23°C or 28°C. Two days after primary local inoculation, upper systemic leaves were infiltrated with *Pst* DC3000 (OD600 = 0.001), and plants were incubated again at their respective temperatures (23°C or 28°C). Bacterial numbers (upper panels) and symptom expression photos (lower panels) were taken at 3 days post-inoculation (dpi) of systemic tissues. Data show the mean log CFU *Pst* DC3000/cm^2^ (± S.D.) and individual points (n=16 from 4 independent experiments) analyzed with two-way ANOVA and Tukey’s Multiple Comparisons test. Statistical differences of means are denoted by different letters. (C) Transcriptome analysis using SAR+ genes from Hartmann et al. (2018) interfaced with temperature-regulated genes from Kim et al. (2022). The number of SAR+ genes that are downregulated and upregulated at elevated temperature are shown (fold change cutoff > 2). The full gene lists are shown in Supplementary Data File 1. (D) Gene Ontology (GO) enrichment analyses of the warm temperature-upregulated SAR+ genes (150) and warm temperature-downregulated SAR+ genes (1037). The full GO results are included in Supplementary Data File 2. (E) Pathogen-induced RPKM values of select plant immunity-associated genes that were downregulated at elevated temperature. For C and E, the heat maps show the log_2_-transformed value of the *Pst*/mock RPKM values at each temperature condition.

In addition to virulent pathogens, avirulent pathogens that induce local ETI can also induce SAR (Cameron et al., 1994; Zeier, 2021). We therefore tested whether SAR activated by the avirulent strain *Pst* DC3000/AvrRpt2, which activates RPS2-dependent immunity in Col-0 plants (Kunkel et al., 1993; Bent et al., 1994), is also impacted by warm conditions. As expected, SAR was induced at normal temperature (23°C) after primary *Pst* DC3000/AvrRpt2 infection (Figure 1B). Strikingly, systemic *Pst* DC3000 levels remained similar compared to mock treatment at elevated temperature (28°C), indicating that even ETI-induced SAR protection is also negatively affected in *Arabidopsis* Col-0 plants at higher temperature.

To further understand global SAR immune signaling, we analyzed a previously generated *Arabidopsis* SAR transcriptome after virulent *P. syringae* infection (Hartmann et al., 2018). We interfaced the SAR-induced (SAR+) genes from that study with our previously published temperature-regulated transcriptome (Kim et al., 2022) and categorized *Arabidopsis* SAR+ genes into downregulated or upregulated genes at elevated temperature. As shown in Figure 1C and Supplementary Data File 1, 1037 SAR+ genes were downregulated, while 151 SAR+ genes were upregulated at elevated temperatures. Gene ontology enrichment analyses (Figure 1D and Supplementary Data File 2) revealed that SAR genes that were upregulated at elevated temperature are associated with toxin catabolism, glutathione metabolism, tryptophan biosynthesis and responses to stress (e.g. hypoxia). In contrast, warm temperature-downregulated SAR genes were predominantly related to defence responses and plant immunity (Figure 1D).

As shown in Figure 1E, these heat-suppressed genes include central components of pattern-triggered immunity (e.g. *FRK1, BIR1, SOBIR1, SERK4*, various PRR genes), effector-triggered immunity (e.g. *EDS1, PAD4, SAG101, NDR1, NSL1, NRG1.2, NRG1.3, ADR1-L1*, various NLR genes), SA pathway (e.g. *ICS1, EDS5, PBS3, NPR3, NPR4, TGA3, PR* genes), NHP pathway (e.g. *ALD1, SARD4, UGT76B1*), calcium signaling (e.g. *ACD6, CNGCs, GLRs, CAMs, CMLs, CPKs*), immune-associated kinases (e.g. MAP kinase and RLCK genes) and defence-related transcription factors (e.g. *WRKYs, NACs, CBP60g, SARD1*). Our transcriptome meta-analyses demonstrated that a significant majority of temperature-regulated SAR genes exhibit downregulated expression at elevated temperature, consistent with loss of systemic protection against secondary infection at 28°C (Figures 1A-B). Overall, our collective results indicate that *Arabidopsis* SAR induced by both virulent and avirulent pathogens are negatively impacted by warm temperatures.

### Temperature regulation of SAR-associated signals

To determine the mechanism of how temperature regulates systemic immunity, we re-examined our temperature-regulated *Arabidopsis* immune transcriptome dataset (Kim et al., 2022) for differential expression of genes involved in various SAR-activating signals. As shown in Figure S1, these included genes important for NHP biosynthesis, nitric oxide generation, reactive oxygen species burst, azelaic acid production, AzA signaling, glycerol-3-phosphate biosynthesis, NADP^+^ generation and SAR-associated auxin response factors (*ARFs*) (Yu et al., 2013; Chen et al., 2018; Hartmann et al., 2018; Shine et al., 2022; Li et al., 2023; Cao et al., 2024).

We found SAR metabolite-associated genes that were either insensitive to temperature change or differentially regulated by elevated temperature. Temperature-insensitive SAR genes included those associated with ROS production (*RBOHD, RBOHF*), NADP^+^ biosynthesis (*FIN4*), auxin response (*ARF1, ARF3, ARF4*), NO generation (*NIA2*, *NOA1*), AzA biosynthesis (*MGD1, DGD1*) and an AzA signaling gene (*DIR1-LIKE*) (Figure S1). On the other hand, warm temperature-downregulated genes included all Pip and NHP biosynthetic genes (*ALD1, SARD4, FMO1*), two AzA signaling gene (*AZI1, EARLI1*) and a G3P gene (*GLI1*). Finally, a second G3P gene (*GLY1*) was actually upregulated at elevated temperature (Figure S1). Collectively, transcriptome analyses show that warming temperature conditions differentially regulate the expression of diverse SAR-associated genes.

### The *N*-hydroxypipecolic acid pathway is downregulated by elevated temperature

Because the Pip-NHP pathway acts upstream of NO-ROS amplification and AzA-G3P signalling during SAR (Wang et al., 2018a), we ultimately focused on the NHP biosynthetic pathway (Figure 2A) as the potentially rate-limiting node in temperature-sensitive systemic immunity. To validate the transcriptome results for the NHP pathway at a narrow temperature range (23-28°C), we measured expression levels of NHP biosynthetic genes (Hartmann et al., 2018; Chen et al., 2018) using RT-qPCR analyses. As shown in Figure 2B-C, transcript levels of *ALD1* and *FMO1* in local (primary) leaves of *Arabidopsis* Col-0 plants at 1 day post-infection with *Pst* DC3000 were lower at 28°C than at 23°C. In agreement, pathogen-induced levels of NHP and its precursor metabolite Pip were also lower at the warmer temperature (Figure 2D-E). These results indicate that elevated temperature negatively impacts the NHP biosynthetic pathway, which is associated with loss of SAR at increasing temperatures.

**Figure 2.**
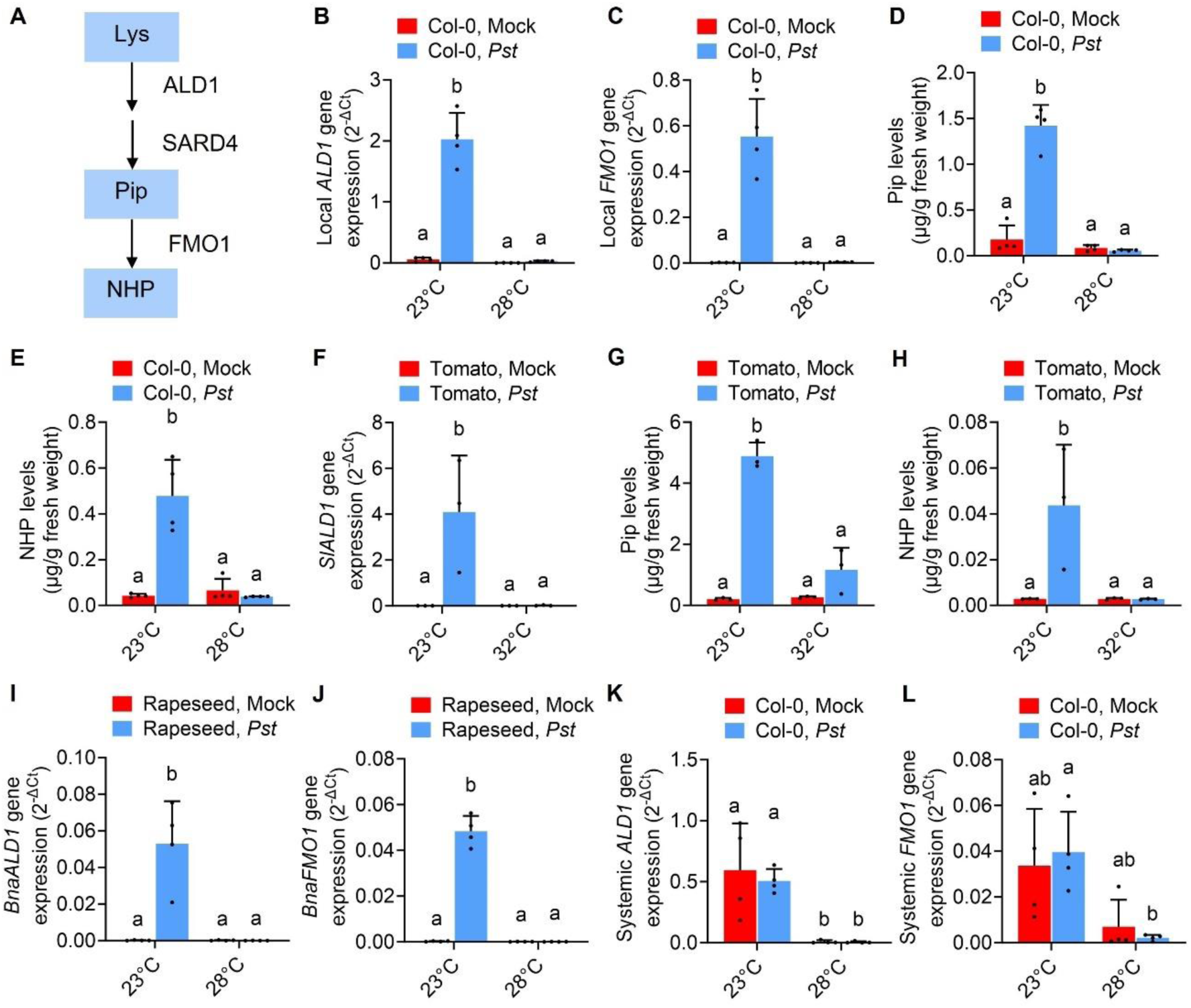
Plant NHP biosynthetic gene expression and NHP production are suppressed at elevated temperature. (A) Simplified diagram of Pip and NHP biosynthesis. (B-E) Leaves of four-week-old *Arabidopsis* Col-0 plants were infiltrated with 0.25 mM MgCl_2_ (mock) or *Pst* DC3000 (OD600 = 0.001). Plants in were then incubated at either 23°C or 28°C. (B) *ALD1* and (C) *FMO1* transcript levels, as well as (D) Pip and (E) NHP levels of pathogen-inoculated tissues were measured at 1 day post-inoculation (dpi). (F-H) Leaves of four-week-old tomato cultivar Castlemart (F) or Moneymaker plants (G-H) were infiltrated with 0.25 mM MgCl_2_ (mock) or *Pst* DC3000 (OD600 = 0.001). (F) *SlALD1* transcript levels, (G) Pip and (H) NHP levels of pathogen-inoculated tissues were measured at 1 dpi. (I-J) Leaves of 4- to 5-week-old rapeseed cultivar Westar plants were infiltrated with 0.25 mM MgCl_2_ (mock) or *Pst* DC3000 (OD600 = 0.0001). (I) *BnaALD1* and (J) *BnaFMO1* transcript levels of pathogen-inoculated tissues were measured at 1 dpi. (K-L) Lower leaves of four-week-old *Arabidopsis* Col-0 plants were infiltrated with 0.25 mM MgCl_2_ (mock) or *Pst* DC3000 (OD600 = 0.02). Plants in were then incubated at either 23°C or 28°C. (K) *ALD1* and (L) *FMO1* transcript levels in upper systemic tissues were measured at 2 dpi. Data show the means (± S.D.) and individual points (n=3 to 4) analyzed with two-way ANOVA and Tukey’s Multiple Comparisons test. Statistical differences of means are denoted by different letters. Experiments were performed at least two times with reproducible results.

Similar trends in warm temperature-suppression of NHP biosynthetic gene expression and/or NHP metabolite levels were observed in tomato plants (Figures 2F-H) and rapeseed plants (Figures 2I-J). Furthermore, as shown in Figures 2K-L, systemic (uninfected) leaves of *Arabidopsis* plants at 2 dpi exhibited lower *ALD1* and *FMO1* gene expression levels at 28°C compared to those at 23°C. Finally, we also observed loss of *Arabidopsis* SAR at intermediate temperatures 24.5°C and 26.5°C (Figure S2), which was associated with loss of pathogen-induced *ALD1* gene expression in *Arabidopsis* plants (Figure S3). Considering that the plant species tested in this study belong to phylogenetically separate families (Brassicaceae and Solanaceae), our findings highlight the remarkable conservation of heat-mediated suppression of NHP biosynthesis in diverse plant lineages.

Because SA is functionally linked and mutually amplified with Pip and NHP during plant systemic immunity (Hartmann and Zeier, 2019; Huang et al., 2020; Zeier, 2021; Shields et al., 2022), we also measured SA biosynthetic gene expression (Figure 3A) in upper, systemic leaves after primary pathogen challenge. As shown in Figures 3B and 3C, systemic expression of the SA biosynthetic gene *ICS1* (Wildermuth et al., 2001) was induced at 2 days after virulent *Pst* DC3000 or avirulent *Pst* DC3000/AvrRpt2 inoculation at 23°C. However, *ICS1* transcript levels in systemic tissues were comparable between mock and pathogen treatments at 28°C (Figures 3A-B), suggesting that elevated temperature downregulates SA accumulation systemically. *ICS1* transcript levels in pathogen-induced plant tissues were also downregulated at elevated temperature (Figure 3D), consistent with previous studies (Huot et al., 2017; Kim et al., 2022).

**Figure 3.**
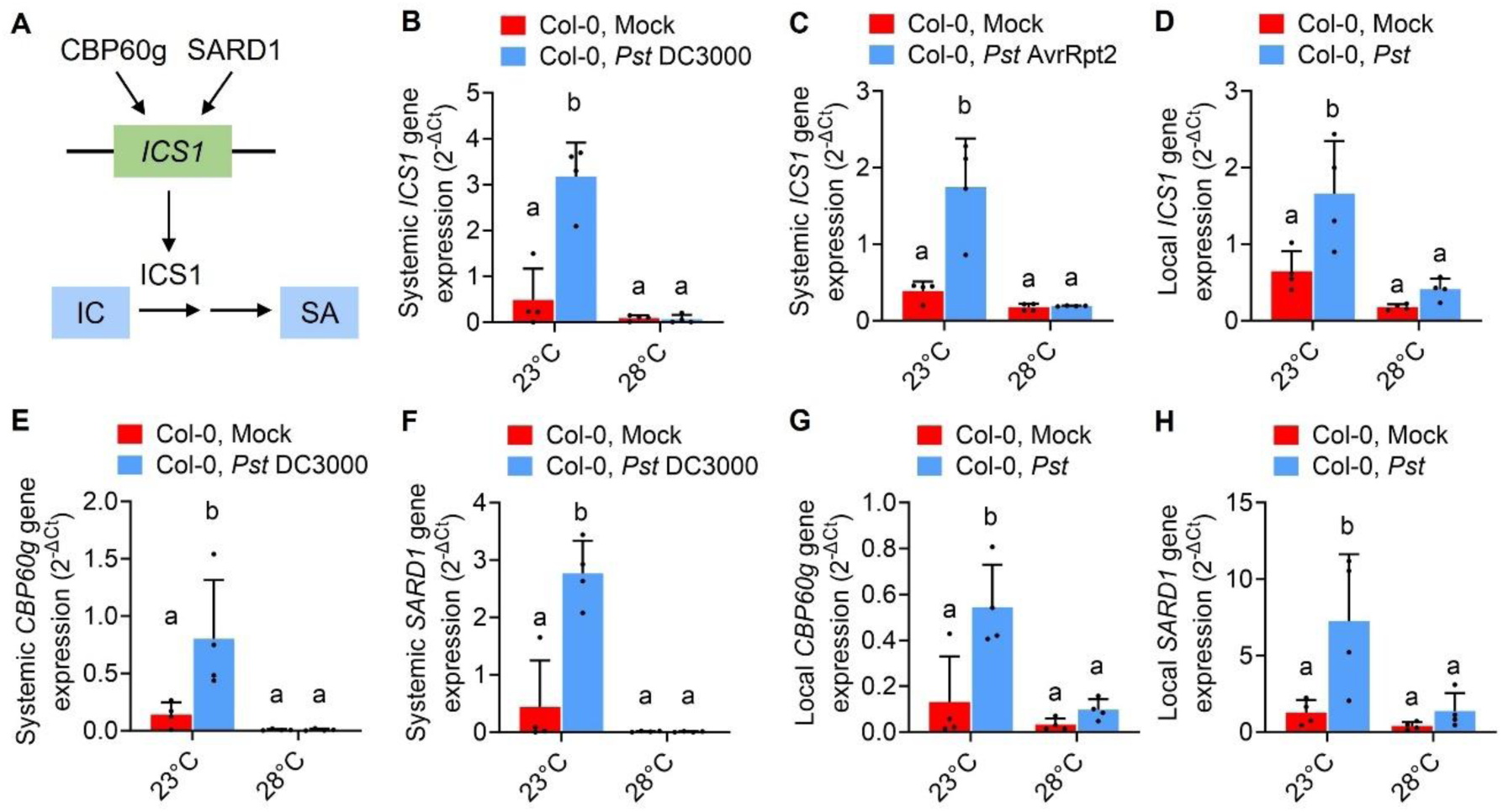
Systemic and local SA biosynthetic gene expression are suppressed at elevated temperature. (A) Simplified diagram of ICS1-mediated SA biosynthesis controlled by master transcription factors CBP60g and SARD1. (B-C) Lower leaves of four-week-old *Arabidopsis* Col-0 plants were infiltrated with 0.25 mM MgCl_2_ (mock), *Pst* DC3000 (OD600 = 0.02) or *Pst* DC3000 AvrRpt2 (OD600=0.02). Plants were then incubated at either 23°C or 28°C. *ICS1* transcript levels were measured in systemic tissues using RT-qPCR at 2 days post-inoculation (dpi) with *Pst* DC3000 (B) or *Pst* DC3000 AvrRpt2 (C), respectively. (D) Leaves of four-week-old *Arabidopsis* Col-0 plants were infiltrated with 0.25 mM MgCl_2_ (mock) or *Pst* DC3000 (OD600 = 0.001) and then incubated at either 23°C or 28°C. *ICS1* transcript levels were measured in pathogen-infected tissues using RT-qPCR at 1 dpi. (E-F) *CBP60g* and *SARD1* transcript levels were measured in systemic tissues at 2 dpi with *Pst* DC3000. (G-H) *CBP60g* and *SARD1* transcript levels were measured in pathogen-infected tissues at 1 dpi with *Pst* DC3000. Data show the means (± S.D.) and individual points (n=4) analyzed with two-way ANOVA and Tukey’s Multiple Comparisons test. Statistical differences of means are denoted by different letters. Experiments were performed at least two times with reproducible results.

Immune-induced *ICS1* gene expression is redundantly regulated by the master transcription factors CBP60g and SARD1 (Wang et al., 2009; Zhang et al., 2010; Wang et al., 2011; Figure 3A). In addition, CBP60g and SARD1 also directly regulate NHP biosynthetic gene expression (*ALD1, SARD4, FMO1*) after pathogen infection (Sun et al., 2015; Sun et al., 2018). Therefore, we also quantified *CBP60g* and *SARD1* expression by RT-qPCR analyses. Consistent with *ICS1* suppression under warm conditions, *CBP60g* and *SARD1* transcript levels were induced at 23°C in local and systemic tissues but this immune induction was lost at 28°C (Figures 3E-H). Altogether, these results demonstrate that the SA biosynthetic pathway is suppressed at higher temperature similarly as NHP. This suggests that mutual amplification of these two important SAR-associated metabolites (SA and NHP) is negatively affected when temperatures increase.

### Exogenous NHP restores *Arabidopsis* immune priming at elevated temperature

Having observed that NHP levels were suppressed at elevated temperature, we hypothesized that exogenous supplementation with NHP may restore disease protection at 28°C if NHP production is the rate-limiting step at elevated temperature. We infiltrated leaves of *Arabidopsis* plants with mock or 1 mM NHP and then infected the same leaves with *Pst* DC3000. As shown in Figure 4A-B, disease symptoms and pathogen levels were reduced after NHP treatment compared to mock treatment at 23°C (∼13-fold reduction in *Pst* pathogen levels), expectedly indicating that NHP is sufficient to induce immune priming. Remarkably, NHP-induced disease protection was maintained at 28°C with a ∼27-fold reduction in *Pst* pathogen levels) (Figure 4A-B), which confirms that NHP production is a rate-limiting step in SAR at elevated temperature.

**Figure 4.**
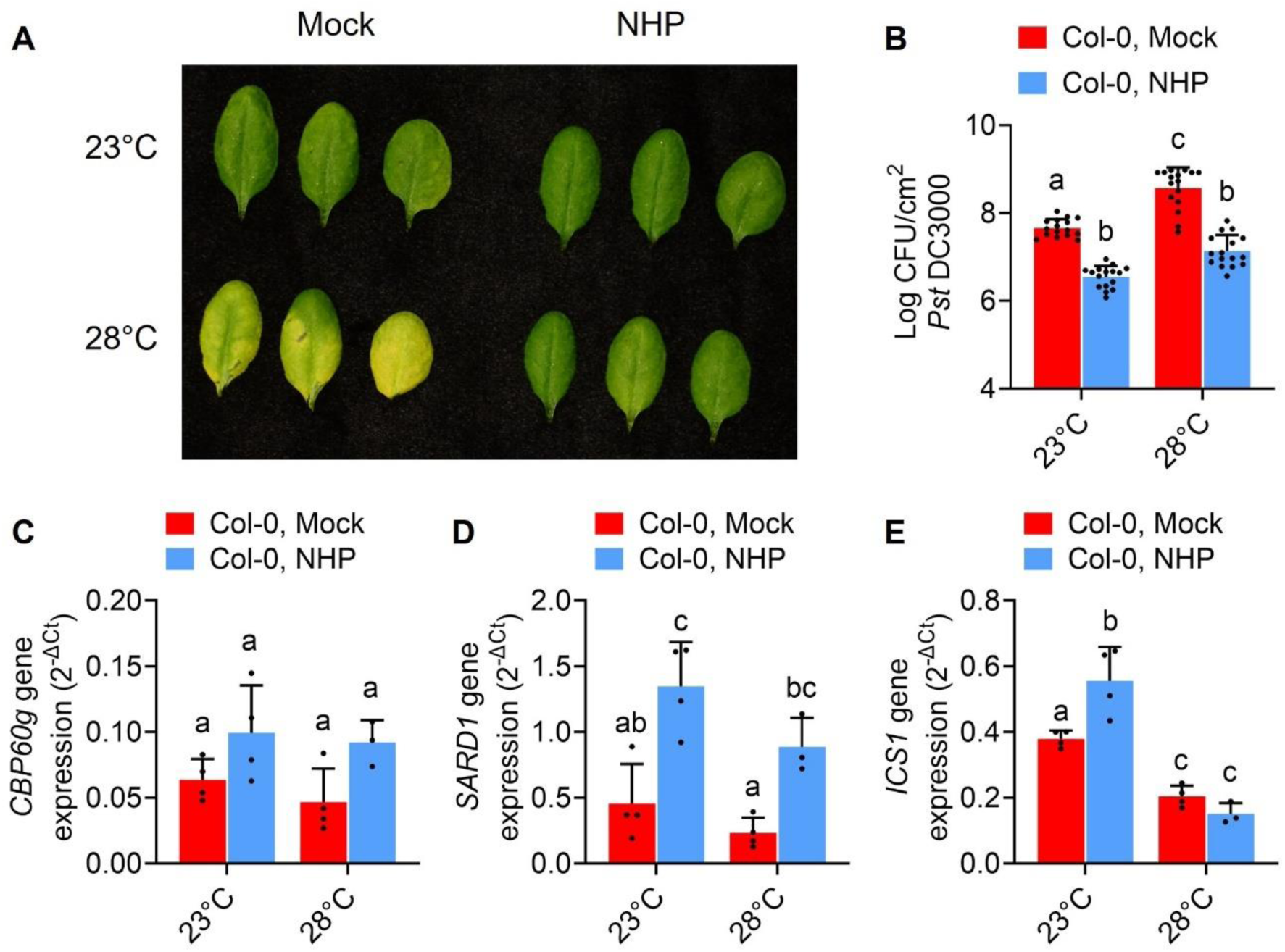
Exogenous NHP treatment restores *Arabidopsis* immune priming at warm temperature. Four-week-old *Arabidopsis* Col-0 plants were treated with mock or 1mM NHP solution. Plants were then incubated at either 23°C or 28°C. (A-B) Two days after NHP treatment, leaves were infiltrated with *Pst* DC3000 (OD600 = 0.001), and plants were incubated again at their respective temperatures (23°C or 28°C). Disease symptoms (A) and bacterial numbers (B) were quantified at 3 days post-inoculation (dpi). Data show the mean log CFU *Pst* DC3000/mL (± S.D.) and individual points (n=16 from 4 independent experiments) analyzed with two-way ANOVA and Tukey’s Multiple Comparisons test. (C-E) NHP-treated leaves were collected after 24h. *CBP60g* (C), *SARD1* (D) and *ICS1* (E) transcript levels were measured with RT-qPCR. Data show the means (± S.D.) and individual points (n=4) analyzed with two-way ANOVA and Tukey’s Multiple Comparisons test. Statistical differences of means are denoted by different letter. Experiments were performed three times with reproducible results.

Because NHP was sufficient to induce immune priming at both temperatures in *Arabidopsis* plants, we investigated whether NHP-mediated immune resilience at warm temperature is due to restored expression of rate-limiting defence genes, such as those involved in SA biosynthesis. As shown in Figure 4C-D, NHP was capable of significantly inducing *SARD1* at both temperatures but not *CBP60g*. Surprisingly, NHP-induced *ICS1* gene expression remained suppressed at elevated temperature (Figure 4E) in spite of the sustained effectiveness of NHP in protecting against *Pst* challenge at 28°C. Similar to the results with NHP-induced disease protection, we also observed reduced pathogen levels after Pip treatment at both 23°C and 28°C (Figure S4). Pip led to a ∼3-fold reduction in *Pst* pathogen levels after leaf infiltration at both temperatures and a 3-4-fold decrease in pathogen load after root treatment (Figure S4). Collectively, our results indicate that the temperature-suppression of the Pip-NHP pathway is governed at the level of Pip and NHP biosynthesis. These findings also indicate that NHP-mediated disease protection was preserved at warmer conditions even without a corresponding *ICS1* gene induction by NHP, suggesting ICS1-independent mechanisms of NHP-mediated immune resilience.

### *CBP60g* and *SARD1* control temperature-vulnerability of the NHP-SAR pathway

In addition to chemical supplementation, we next investigated if we could restore systemic immune responses genetically at warm temperatures. We recently showed that CBP60g and its functionally redundant paralog SARD1 control the temperature-sensitivity of plant basal resistance and pathogen-induced SA biosynthesis (Kim et al., 2022). We therefore investigated whether CBP60g and SARD1 also control temperature-sensitive systemic defence responses. As shown in Figure 3E-H, *CBP60g* and *SARD1* transcript levels are induced locally and systemically after *Pst* DC3000 challenge at 23°C, but this induction is lost at 28°C. These suggest that the loss of systemic immunity and NHP biosynthesis in *Arabidopsis* plants at elevated temperature could be due to significantly decreased *CBP60g* and *SARD1* gene expression.

To show a causative role for *CBP60g/SARD1* downregulation with NHP immune pathway suppression, we used plants constitutively expressing *CBP60g* (*35S::CBP60g*) or *SARD1* (*35S::SARD1*). As shown in Figures 5A-C, *35S::CBP60g* lines restored NHP biosynthetic gene expression (*ALD1, FMO1*) and levels of the NHP precursor Pip. Consistent with the previously defined functional redundancy of CBP60g and SARD1 (Zhang et al., 2010; Wang et al., 2011; Sun et al., 2015; Kim et al., 2022), constitutive *SARD1* expression in *35S::SARD1* plants also led to restored expression of *ALD1* and *FMO1* at warm temperature (Figure 5D-E).

**Figure 5.**
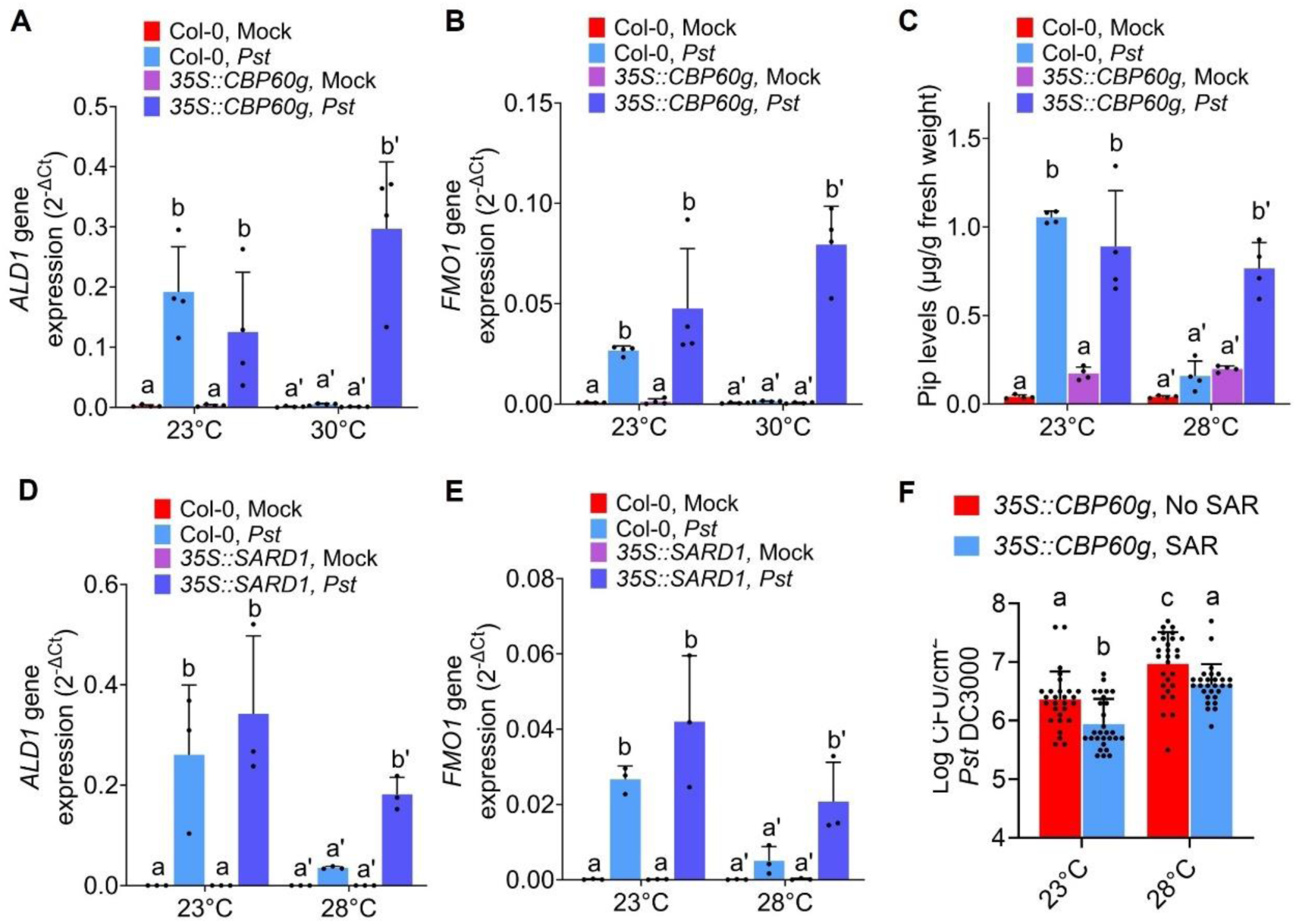
CBP60g and SARD1 control the temperature-vulnerability of NHP-mediated immunity. Leaves of four-week-old *Arabidopsis* Col-0 and *35S::CBP60g* (A-C) or *35S::SARD1* plants (D-E) were infiltrated with 0.25 mM MgCl_2_ (mock) or *Pst* DC3000 (OD600 = 0.001). Plants were then incubated at either 23°C or 28°C-30°C. (A) *ALD1* and (B) *FMO1* transcript levels were measured using RT-qPCR at 1 day post-inoculation (dpi) of pathogen-inoculated tissues of Col-0 and *35S::CBP60g* plants. (C) Pip levels were measured at 1 dpi in the same tissues. (D) *ALD1* and (E) *FMO1* transcript levels were measured at 1 day post-inoculation (dpi) of pathogen-inoculated tissues of Col-0 and *35S::SARD1* plants. Data show the means (± S.D.) and individual points (n=4 in A-C; n=3 in D-E), with experiments performed three times with reproducible results. (F) Four-week-old *Arabidopsis 35S::CBP60g* plant leaves were infiltrated with 0.25 mM MgCl_2_ (mock) and *Pst* DC3000/AvrRpt2 (OD600 = 0.02). Plants were then incubated at either 23°C or 28°C. Two days after primary local inoculation, upper systemic leaves were infiltrated with *Pst* DC3000 (OD600 = 0.001), and plants were incubated again at their respective temperatures (23°C or 28°C). Bacterial numbers were taken at 3 days post-inoculation (dpi) of systemic tissues. Data show the mean log CFU *Pst* DC3000/cm^2^ (± S.D.) and individual points (n=28 from 7 independent experiments) analyzed with two-way ANOVA and Tukey’s Multiple Comparisons test. Statistical differences of means are denoted by different letters.

In terms of the biological relevance of restored NHP biosynthesis with constitutive *CBP60g* expression, we observed effective SAR in *35S::CBP60g* plants at both normal and elevated temperature (Figure 5F), in contrast to wild-type Col-0 plants that lose SAR protection at 28°C (Figure 1A-B). Intriguingly, *35S::SARD1* plants had consistently enhanced disease resistance even without initial SAR priming at both temperatures, which was not accompanied by additional reduction in systemic pathogen levels after SAR priming (Figure S5).Taken together, a major mechanism by which higher temperatures target plant systemic immunity and NHP biosynthesis is through the expression of the master immune transcription factor genes *CBP60g* and *SARD1*.

## Discussion

In this study, we showed that pathogen-induced SAR and the SAR-activating NHP pathway in *Arabidopsis* plants are sensitive to elevated temperatures. Local infection with both virulent or ETI-activating avirulent pathogens triggers SAR at normal temperatures but not at elevated temperatures. We showed that SAR regulation by temperature is caused by temperature-sensitive NHP pathway in local and systemic tissues. SAR-induced NHP biosynthetic gene expression (*ALD1, FMO1*) and levels of both NHP (and its precursor Pip) are downregulated at elevated temperature. Even though multiple SAR signals have been proposed previously (Fu and Dong, 2013; Návarová, et al., 2012; Hartmann et al., 2018; Wang, et al., 2018; Vlot et al., 2021; Zeier, 2021), we show here that exogenous supplementation with NHP or Pip (locally and systemically) was sufficient to restore SAR at elevated temperature. This demonstrates a causative relationship between temperature-suppressed NHP pathway and SAR. In the future, it would be interesting to disentangle the differential temperature effects on both the initial SAR priming with *Pst* DC3000/AvrRpt2 and the secondary pathogen challenge with *Pst* DC3000.

What is clear is the rate-limiting nature of NHP biosynthesis in conferring immune preparedness in plants, which is a major concern in a changing climate with more sustained heat waves. It is interesting to note that there are both temperature-sensitive (exemplified by *ALD1* and *FMO1* in the NHP pathway) and temperature-insensitive genes associated with generating central SAR signals. These temperature-insensitive SAR nodes include *RBOHD/RBOHF* (for ROS generation), *FIN4* (for NADP^+^ biosynthesis), *ARF1/3/4* (for tasi-ARF signaling), *NIA2/NOA1* (for NO generation), and *MGD1/DGD1* (for AzA biosynthesis). Because NHP has been shown to work upstream of most of these SAR signals (Wang et al., 2018a; Li et al., 2023), these temperature-regulated trends underscore that indispensability of the NHP in conferring robust systemic immunity under changing environmental conditions. Future studies can then address the individual contributions of these SAR signals downstream of NHP (e.g. NADP^+^, ROS, AzA and G3P) in conferring systemic disease resistance under variations in temperatures and other climatic factors.

Because NHP primes systemic SA biosynthesis and immunity (Yildiz et al., 2021; Zeier, 2021; Li et al., 2023), we also observed that systemic SA biosynthetic gene expression is downregulated by elevated temperature. SA induction is important for SAR establishment since SA induction-deficient *sid2* and SA-insensitive *npr1* mutants also have abolished SAR (Fu and Dong, 2013; Huang et al., 2020; Peng et al., 2021). Temperature-suppressed systemic SA pathway agrees with our previous study showing a negative impact of higher temperatures on local pathogen-induced SA biosynthesis and basal resistance (Huot et al., 2017; Kim et al., 2022). Remarkably, we found that NHP or Pip supplementation alone can protect against *Pst* pathogen challenge at elevated temperature, suggesting that Pip-NHP signaling is sufficient in inducing SAR. It is interesting to note that NHP-mediated disease protection at warmer conditions is associated with partial but not full restoration of the SA biosynthetic pathway. While *CBP60g* and *SARD1* gene expression remain temperature-resilient after NHP treatment, NHP only induces *ICS1* transcript levels at normal temperature but not at elevated temperature. These suggest *ICS1*-independent mechanisms of plant immune resilience to climate warming, which has been shown in certain natural accession of *A. thaliana* plants (Rossi et al., 2024).

A potential mechanism for NHP-mediated immune resilience to changing temperatures can be gleaned from previously demonstrated roles of NHP in inducing MAPK activation (Wang et al., 2018b) and/or *EDS1* expression (Nair et al., 2021). Because sustained MAPK or EDS1 activation can compensate for lack of SA induction at ambient temperature conditions (Venugopal et al., 2009; Tsuda et al., 2013), this could explain why NHP-induced disease resistance is preserved at warm temperature. Alternatively, basal SA levels that remain at elevated temperature, without a corresponding *ICS1* induction, may be sufficient for NHP-mediated immune priming. This possibility agrees with results in SA induction-deficient mutant *sid2*, which can still induce immune gene expression (like *SARD1* and *FMO1*) after NHP treatment (Nair et al., 2021).

We previously showed that CBP60g and SARD1 controls the temperature-vulnerability of local pathogen-induced SA biosynthesis (Kim et al., 2022). Partially redundant paralogs CBP60g and SARD1 are master transcription factors that directly target the promoters of numerous immunity-related genes (Wang et al., 2009; Zhang et al., 2010; Wang et al., 2011; Sun et al., 2015, Sun et al., 2018), including those important for SA biosynthesis (like *ICS1*) and NHP biosynthesis (like *ALD1* and *FMO1*). In this study, we further demonstrate that CBP60g and SARD1 control the temperature-regulation of NHP biosynthesis and systemic immunity. Expression of *CBP60g* and *SARD1* in both local and systemic tissues are downregulated at elevated temperature, which is associated with suppressed NHP pathway. Restoring organism-wide *CBP60g* or *SARD1* expression using constitutively expressing *35S::CBP60g* or *35S::SARD1* plants rescues NHP biosynthetic gene expression and Pip levels under warming conditions.

In contrast to *35S::CBP60g* plants which exhibits SAR at both 23°C and 28°C, *35S::SARD1* plants seem to already exhibit primed immunity even without SAR induction by initial pathogen infection. This could be due to differences in *CBP60g* and *SARD1* gene or protein regulation. While the CBP60g protein requires interaction with calcium-activated calmodulin protein, SARD1 does not require calmodulin (Wang et al., 2009; Zhang et al., 2010). *SARD1* is also considered a late pathogen-responsive gene, but *CBP60g* is an early immune-elicited gene in terms of expression profiles (Wang et al., 2011). Finally, overexpression of *SARD1* in a previous study has already been shown to exhibit constitutive SAR phenotypes at ambient temperature (Zhang et al., 2010). Altogether, while CBP60g and SARD1 redundantly control the temperature-regulation of NHP biosynthesis in *Arabidopsis*, their differential and potentially distinct functions in plant systemic immunity still need to be further unravelled in future studies.

Overall, changing climatic factors like elevated temperature have a broad and significant impact on the plant immune system, not only at local sites of infection (PTI, ETI, SA) but also on systemic immune priming in distal sites via the central SAR metabolite NHP. In this study, we determined that elevated temperature regulates SAR by influencing the NHP pathway. We further demonstrated that this temperature-regulation of NHP-mediated SAR is also controlled by the temperature-sensitive master regulators CBP60g and SARD1, reminiscent of their roles in local immune responses (Kim et al., 2022). Not only does temperature affect the plant’s ability to directly defend against pathogen attacks but also the plant’s immune preparedness for future infections. Our discoveries advance our understanding of how the plant immune landscape is regulated by a changing environment. This foundational mechanistic knowledge of the plant disease triangle is important to inform strategies in mitigating the negative impacts of warming temperatures on plant health and to provide a molecular roadmap towards engineering climate-resilient plants.

## Materials and Methods

### Plant materials and growth conditions

*Arabidopsis thaliana* Columbia-0 (Col-0), *35S::CBP60g* (Wan et al., 2012) and *35S::SARD1* seeds (Kim et al., 2022) were surface-sterilized with 70% ethanol for 10 minutes and rinsed with autoclaved water. Seeds were suspended in sterile 0.1% agarose and incubated in cold conditions (4°C) for three days. Seeds were sown into pots containing autoclaved soil mix – one-part Promix-PGX soil (Plant Products, Ancaster, Ontario), one-part Turface (Turface Athletics, Buffalo Grove, IL), and one-part Vermiculite Prolite (Therm-O-Rock, New Eagle, PA) – supplemented with 100mL of Miracle-Gro solution (The Scotts Company, Mississauga, ON). Tomato (*Solanum lycopersicum*) cultivar Castlemart and Moneymaker plants were grown as described previously (Shivnauth et al., 2023). Rapeseed (*Brassica napus*) cultivar Westar plants were grown as described previously (Kim et al., 2022). Plants were grown in environmentally controlled chambers at 23°C with 60% relative humidity and 12h light [100 ± 20 µmol m^−2^ s^−1^]/12 dark cycle based on established procedures (Huot et al., 2017; Kim et al., 2022; Rossi et al., 2024).

### Plant systemic immunity and SAR assays

Four-week-old *Arabidopsis* plants were covered with plastic domes to increase humidity and open the stomata 24 hours before infiltration, Plants were infiltrated with 0.25 mM MgCl_2_ (mock), *Pst* DC3000 (OD600=0.02) or *Pst* DC3000/AvrRpt2 (OD600=0.02) in their lower leaves (Xin et al., 2013; Xin et al., 2018). Following infiltration, plants were further incubated in environmentally growth chambers at either 23°C or 28°C with identical relative humidity (60%) and lighting conditions (12h light/12 dark). For SAR disease assays, upper systemic leaves were or infiltrated with *Pst* DC3000 (OD600 = 0.001). Bacterial levels were quantified 3 days after systemic infiltration based on a previously published protocol (Huot et al., 2017; Kim et al., 2022; Rossi et al., 2024). Briefly, in planta bacterial extracts were plated on rifampicin-containing LM media and log colony forming units (CFUs) cm^-2^ were calculated.

### Pip treatment and protection assay

Four-week-old *Arabidopsis* plants were infiltrated with mock solution or 1 mM Pip (Sigma) in their lower leaves. In parallel to direct Pip infiltration into leaves, plants were irrigated with 1 mL of mock solution or 1 mM Pip (Sigma) for root inoculations. Following infiltration or irrigation, plants were further incubated in environmentally growth chambers at either 23°C or 28°C with identical relative humidity (60%) and lighting conditions (12h light/12 dark). After two days, the Pip-treated leaves (for local Pip infiltration) or upper systemic leaves (for root irrigation) were further infiltrated with *Pst* DC3000 (OD600 = 0.001). Bacterial levels were quantified 3 days after systemic infiltration as stated in the previous section.

### NHP treatment and protection assay

Four-week-old *Arabidopsis* plant leaves were infiltrated with mock solution or 1 mM NHP (MedChemExpress) and then incubated in environmentally growth chambers at either 23°C or 28°C with identical relative humidity (60%) and lighting conditions (12h light/12 dark). After one day, the NHP-treated leaves were collected for gene expression analyses or further infiltrated with *Pst* DC3000 (OD600 = 0.001). In the latter case, bacterial levels were quantified 3 days after systemic infiltration as detailed previously.

### Gene expression analyses

Pathogen-infected leaves and upper systemic leaves at either 23°C or 28°C were harvested at 1 or 2 days after local infection, respectively. NHP-infiltrated leaves at both temperatures were harvested at 1 day after NHP treatment. Tissues were flash-frozen in liquid nitrogen and stored at -80°C before total RNA extraction. Gene expression levels were quantified based on a previously published protocol (Huot et al., 2017; Kim et al., 2022) with slight modifications. RNA was extracted from flash-frozen plant tissues using the Qiagen Plant RNeasy Mini Kit (Qiagen, Toronto, ON) or TRIzol Reagent (Aidlab Biotech, China) according to the manufacturer’s protocol. Resulting cDNA was synthesized using qScript cDNA super mix (Quantabio) or M-MLV reverse transcriptase (RT, Vazyme Biotech, China) based on manufacturers’ recommendations. Real-time quantitative polymerase chain reaction (qPCR) was performed using PowerTrack SYBR Green master mix (Life Technologies) or iTaq™ Universal SYBR® Green Supermix (Bio-Rad, USA) with approximately 1.5-10 ng of template cDNA. Equivalently diluted mRNA without the qScript cDNA mix were used as negative controls. The resulting qPCR mixes were run using the Applied Biosystems QuantStudio3 platform (Life Technologies) or CFX96 Real-Time PCR Detection System (Bio-Rad, USA). The individual Ct values were determined for target genes and the internal control gene: *PP2AA3* for Arabidopsis; *SlACT2* for tomato and *BnaGDI1* for rapeseed (Huot et al., 2017; Kim et al., 2022; Shivnauth et al., 2023). Gene expression values were reported as 2^−ΔCt^, where ΔCt is Ct_target gene_– Ct_internal control gene_. qPCR was carried out with three technical replicates for each biological sample. Primers used for qPCR are shown in Supplementary Table 1.

### Transcriptome meta-analyses

The SAR+ gene list was retrieved from a previously published transcriptome dataset (Hartmann et al., 2018). We then interfaced all SAR+ genes with the temperature-regulated transcriptome dataset from Kim et al. (2022). At each temperature (23°C or 28°C), the fold-change induction after *Pst* infection (*Pst*/mock) was first calculated. The 28°C/23°C *Pst* induction ratios were then determined for all SAR+ genes. Genes were classified as either downregulated or upregulated at elevated temperature based on a fold-change cutoff greater than 2. The two gene clusters (warm-downregulated and warm-upregulated) were then subjected to Gene Ontology (GO) analyses using David (Sherman et al., 2022)

### Pip and NHP metabolite extraction and quantification

Pathogen-infected leaves at 23°C or 28°C were harvested at 1 day after infection. Pip extraction and quantification were performed based on previous reports with slight modifications (Yao et al., 2023; Yao et al., 2024). Approximately 100 mg leaf tissue was frozen and ground in liquid nitrogen. Pip and NHP were simultaneously extracted at 4℃ for 1h in 600 µL ice-cold extraction buffer (80% methanol in water, 0.1 g/L butylated hydroxytoluene). The extraction step was repeated twice, and a 1.2-mL supernatant was speed-dried in a vacuum centrifugal concentrator (Beijing JM Technology). The pellet was resuspended in 240 µL 30% methanol solution. The solution remains undiluted for the quantification of NHP, yet it is diluted tenfold for the quantification of Pip. Pip and NHP levels were quantified using the AB SCIEX QTARP 5500 LC/MS/MS system. Selected ion monitoring (SIM) was conducted in the positive ES channel for Pip (m/z 130.0>84.0) and NHP (m/z 146.1>100.0), which was done using a collision energy of 20V (for Pip) or 14V (for NHP) and a declustering potential of 50V (for Pip) or 40V (for NHP). The instrument control and data acquisition were performed using Analyst 1.6.3 software (AB SCIEX), and data processing was performed using MultiQuant 3.0.2 software (AB SCIEX). Pip and NHP were separated with an ACQUITY XSelect HSS T3 column (2.5 µm, 3.0 x 100 mm, Waters) using the method in Supplementary Table 2. Pip and NHP levels were quantified by calculating the area of each individual peak and comparing it to standard curves. Reported Pip and NHP concentrations were normalized by sample fresh weight (FW) in grams.

## Acknowledgments

We thank the following colleagues for sharing biological materials: G. Howe for Castlemart tomato, X. Meng for Moneymaker tomato, B. Staskawicz for *Pst* DC3000/AvrRpt2, and G. Li and D. Wan for *35S::CBP60g*. We are grateful to the Laurier Faculty of Science (Max Pottier and Gena Braun), Michigan State University (MSU) Growth Chamber Facility, Duke Phytotron and CAS Center for Excellence in Molecular Plant Sciences Metabolomics-Mass Spectrometry facility (Yuanyuan Gao) for instrumentation support. We also thank Castroverde Lab members, Robin Cameron (McMaster University) and Keiko Yoshioka (University of Toronto) for meaningful discussions about our study. This work was supported by research funding from the NSERC Discovery Grant, Canada Foundation for Innovation, Ontario Research Fund, Wilfrid Laurier University Faculty of Science start-up funds and the Youth Program of National Natural Science Foundation of China. Our Laurier Biology research is located on the shared traditional territory of the Neutral, Anishinaabe, and Haudenosaunee peoples.

## Author Contributions

C.D.M.C conceptualized and supervised the study. A.S. performed most of the SAR disease assays and gene expression experiments. L.Y.Y. optimized and performed the Pip and NHP metabolite measurements. C.A.M.R. conducted additional SAR assays and gene expression analyses. P.C.C. did the NHP-mediated disease resistance and RT-qPCR assays. J.H.K. performed gene expression experiments of *35S::SARD1* plants. W.M.A.A.T. and E.M. conducted experimental replicates of the SAR assays. V.S. conducted temperature experiments of tomato plants, extracted RNA and synthesized cDNA. S.L. and T.C. performed gene expression analyses of rapeseed plants. L.Y.Y., P.C.C. and S.L. were supervised by X.F.X., S.Y.H. and T.C., respectively. Everyone analyzed the data. A.S. and C.D.M.C. wrote the paper with input from all authors.

## Competing Interest Statement

The authors do not declare any known conflicts of interest.

## Notes

### Competing Interest Statement

The authors have declared no competing interest.

### Summary of Updates

This revision includes new experiments, additional analyses and an updated Discussion section.

