## Supplementary Information for "Warm temperature suppresses plant systemic acquired resistance by intercepting *N*-hydroxypipecolic acid biosynthesis"

**Table S1.** Table of primers.

| Primer Name | Sequence | Reference |
| --- | --- | --- |
| *PP2AA3_forward* | GGTTACAAGACAAGGTTCACTC | Huot et al., 2017 |
| *PP2AA3_reverse* | CATTCAGGACCAAACTCTTCAG | |
| *ALD1_forward* | TCGCTTGGCCTCAAGGTTT | Kim et al., 2017 |
| *ALD1_reverse* | CCTTAAAGTGAACCCACAAGTATGG | |
| *FMO1_forward* | TCGGTGCTGGTGTTAGCGGA | Kim et al., 2017 |
| *FMO1_reverse* | CGAGGCTTCGAATACGGTCGGG | |
| *ICS1_forward* | ACTTACTAACCAGTCCGAAAGACGA | Huot et al., 2017 |
| *ICS1_reverse* | ACAACAACTCTGTCACATATACCGT | |
| *CBP60g_forward* | TCGTGGACGCCACCACAAACA | Kim et al., 2017 |
| *CBP60g_reverse* | TCAGCGTTCAGCGGCACGAG | |
| *SARD1_forward* | TCGAGTTGGATTCGTAGCCG | Kim et al., 2017 |
| *SARD1_reverse* | TCGCTTCAGTCATCGCTTCA | |
| *SlACT2_forward* | TTCAACACCCCTGCCATGT | Shivnauth et al., 2023 |
| *SlACT2_reverse* | CCACTGGCATAGAGGGAAAGAA | |
| *SlALD1_forward (Solyc11g044840)* | CGGGTTCTAGAAAGGTTGCC | Wang et al., 2021 |
| *SlALD1_reverse (Solyc11g044840)* | CAATCCACCAGCCTGAGCTA | |
| *BnaGDI1_forward* | GAGTCCCTTGCTCGTTTCC | Yang et al., 2014 |
| *BnaGDI1_reverse* | TGGCAGTCTCTCCCTCAGAT | |
| *BnaALD1_forward (BnaA03g38440D)* | GATACAACAGAGCCTATACCGA | This study |
| *BnaALD1_reverse (BnaA03g38440D)* | TTTCCTAAGAACCTTGTCACCT | |
| *BnaFMO1_forward (BnaA08g22130D)* | TAGCAAATCAAGGAGAAGGTGG | This study |
| *BnaFMO1_reverse (BnaA08g22130D)* | GGTAGAGTAGAACAAGAAGAATGG | |

**Table S2.** UPLC Method for simultaneous Pip and NHP quantification.

| **Time (min)** | **Flow rate (mL/min)** | **A%** | **B%** |
|---|---|---|---|
| 0-2 | 0.4 | 99 | 1 |
| 2-5 | 0.4 | Decrease linearly to 70 | Increase linearly to 30 |
| 5-6 | 0.4 | Decrease linearly to 5 | Increase linearly to 95 |
| 6-7 | 0.4 | 5 | 95 |
| 7-7.1 | 0.4 | Increase linearly to 99 | Decrease linearly to 1 |
| 7.1-10 | 0.4 | 99 | 1 |

A: water with 0.1% (vol/vol) formic acid;
B: methanol;
The column oven was set at 40 ˚C, and the injection volume was 1 μL.

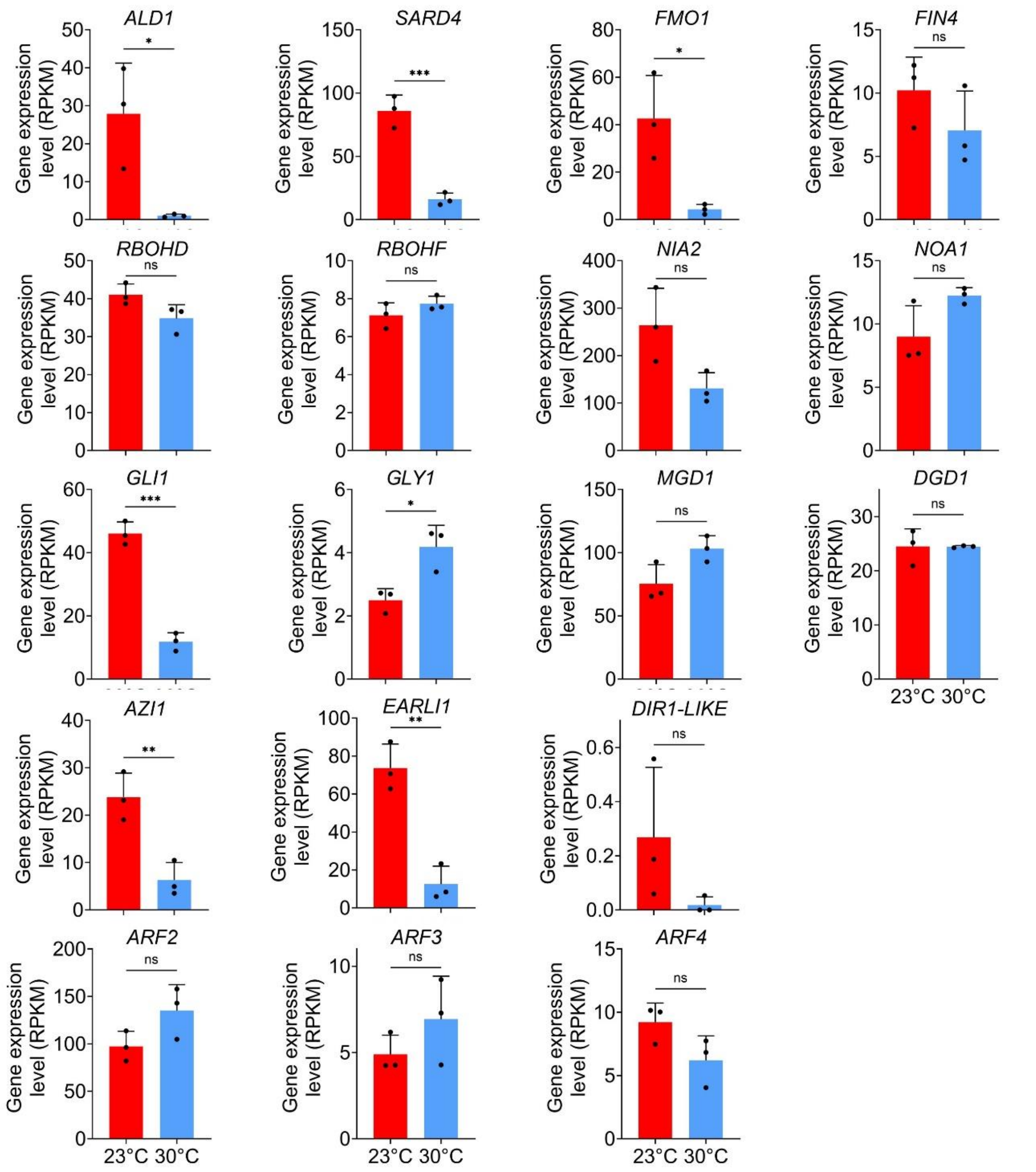


**Figure S1. Transcript levels of SAR-associated genes at normal and elevated temperatures.** The RPKM values of major SAR-associated genes were retrieved from a previous transcriptome (RNA-Seq) study by Kim et al. (2022). The genes of interest are involved in NHP biosynthesis (*ALD1, SARD4, FMO1*), NADP$^+$ generation (*FIN4*), reactive oxygen species burst (*RBOHD, RBOHF*), nitric oxide generation (*NIA2, NOA1*), glycerol-3-phosphate biosynthesis (*GLI1, GLY1*), azelaic acid production (*MGD1, DGD1*), AzA signaling (*AZI1, EARLI1, DIR1-LIKE*) and SAR-associated auxin response factors (*ARF2, ARF3, ARF4*). Data show the means (± S.D.) and individual points (n=3 plants) analyzed with t-test. Statistical differences are denoted by asterisks ("*", $p<0.05$; "**", $p<0.01$, "***", $p<0.001$).

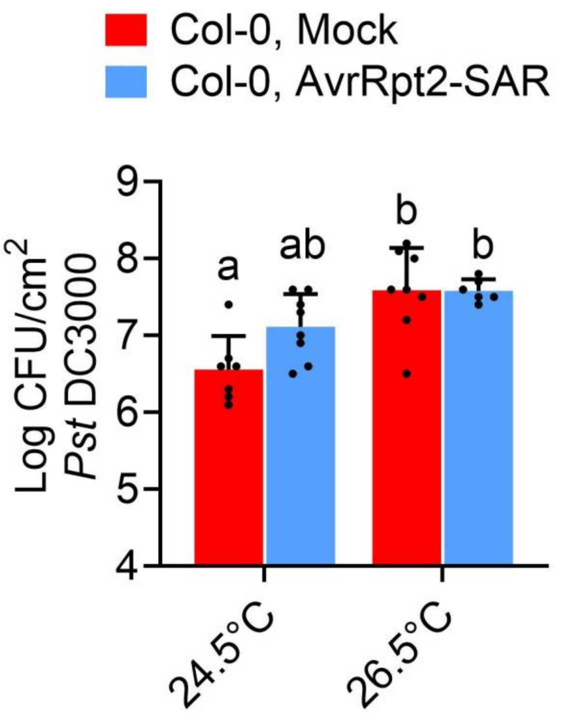


**Figure S2. *Arabidopsis* systemic immune responses at 24.5°C and 26.5°C.** Lower leaves of four-week-old *Arabidopsis* Col-0 plants were infiltrated with 0.25 mM $MgCl_2$ (mock) and *Pst* DC3000/AvrRpt2 (OD600 = 0.02). Plants were then incubated at either 24.5°C or 26.5°C. Two days after primary local inoculation, upper systemic leaves were infiltrated with *Pst* DC3000 (OD600 = 0.001), and plants were incubated again at their respective temperatures (24.5°C or 26.5°C). Bacterial numbers (upper panels) were taken at 3 days post-inoculation (dpi) of systemic tissues. Data show the mean log CFU *Pst* DC3000/cm$^2$ (± S.D.) and individual points (n=7-8 from 2 independent experiments) analyzed with two-way ANOVA and Tukey's Multiple Comparisons test. Statistical differences of means are denoted by different letters.

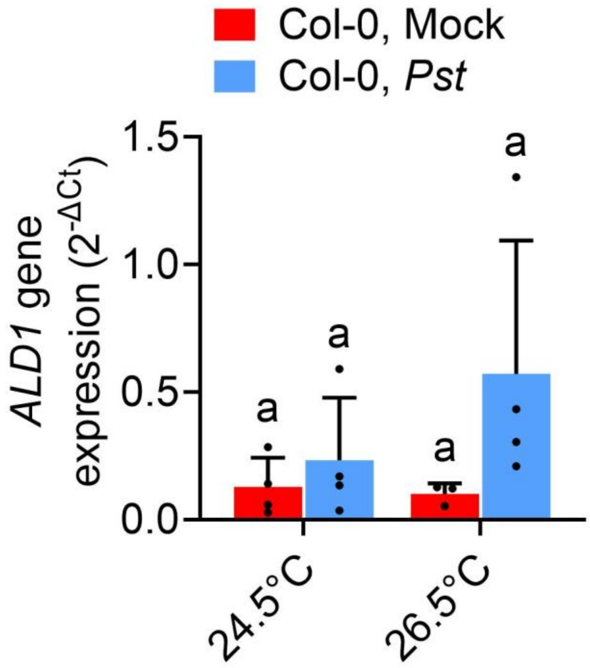


**Figure S3. *ALD1* gene expression in *Arabidopsis* at 24.5°C and 26.5°C.** (A-B) Leaves of four-week-old *Arabidopsis* Col-0 plants were infiltrated with 0.25 mM $MgCl_2$ (mock) or *Pst* DC3000 (OD600 = 0.001). Plants in were then incubated at either 24.5°C or 26.5°C. *ALD1* transcript levels of pathogen-inoculated tissues were measured at 1 day post-inoculation (dpi). Data show the means (± S.D.) and individual points (n=3 to 4) analyzed with two-way ANOVA and Tukey's Multiple Comparisons test. Statistical differences of means are denoted by different letters. Experiments were performed two times with reproducible results.

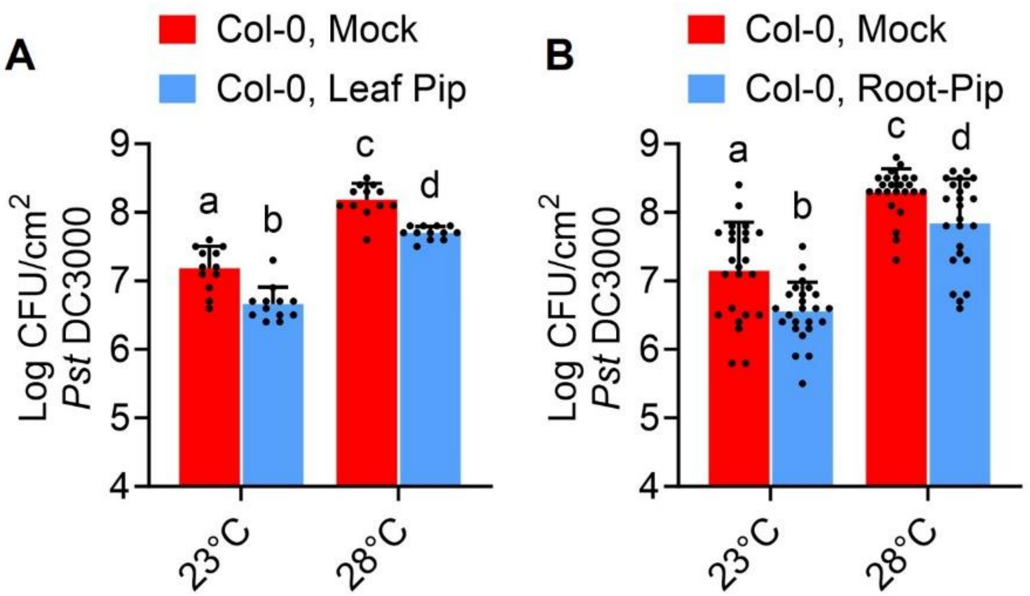


**Figure S4. Exogenous Pip treatment restores *Arabidopsis* immune priming at warm temperature.** Four-week-old *Arabidopsis* Col-0 plants were treated with mock or 1mM Pip solution by leaf-infiltration (A) or root-drenching (B). Plants were then incubated at either 23°C or 28°C. Two days after Pip treatment, leaves were infiltrated with *Pst* DC3000 (OD600 = 0.001), and plants were incubated again at their respective temperatures (23°C or 28°C). Bacterial numbers were quantified at 3 days post-inoculation (dpi). Data show the mean log CFU *Pst* DC3000/mL (± S.D.) and individual points (n=12 from 3 independent experiments in A; n=24 from 6 independent experiments in B) analyzed with two-way ANOVA and Tukey's Multiple Comparisons test. Statistical differences of means are denoted by different letters.

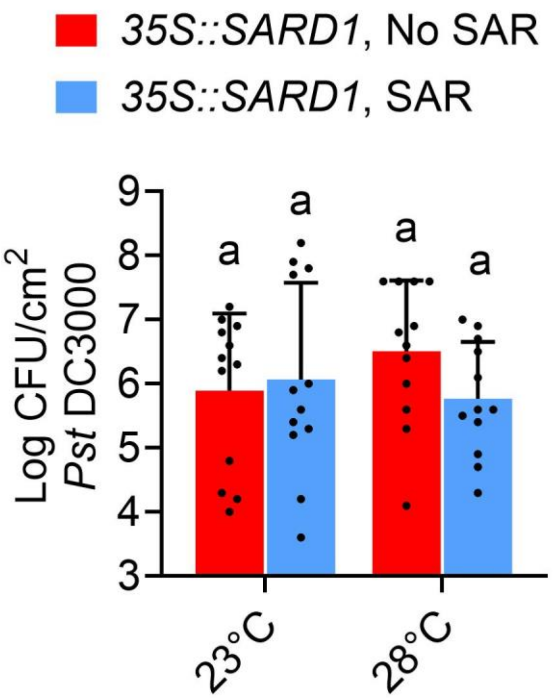


**Figure S5. Systemic acquired resistance phenotypes of *35S::SARD1* plants at normal and elevated temperatures.** Leaves of four-week-old *Arabidopsis 35S::SARD1* plants were infiltrated with 0.25 mM $MgCl_2$ (mock) and *Pst* DC3000/AvrRpt2 (OD600 = 0.02). Plants were then incubated at either 23°C or 28°C. Two days after primary local inoculation, upper systemic leaves were infiltrated with *Pst* DC3000 (OD600 = 0.001), and plants were incubated again at their respective temperatures (23°C or 28°C). Bacterial numbers were taken at 3 days post-inoculation (dpi) of systemic tissues. Data show the mean log CFU *Pst* DC3000/cm$^2$ (± S.D.) and individual points (n=12 from 3 independent experiments) analyzed with two-way ANOVA and Tukey's Multiple Comparisons test. Statistical differences of means are denoted by different letters.
